# Redundancy, feedback, and robustness in the *Arabidopsis thaliana BZR/BEH* gene family

**DOI:** 10.1101/053447

**Authors:** Jennifer Lachowiec, G. Alex Mason, Karla Schultz, Christine Queitsch

## Abstract

Organismal development is remarkably robust, tolerating stochastic errors to produce consistent, so-called canalized adult phenotypes. The mechanistic underpinnings of developmental robustness are poorly understood, but recent studies implicate certain features of genetic networks such as functional redundancy, connectivity, and feedback. Here, we examine the *BRZ/BEH* gene family, whose function is crucial for embryonic stem development in the plant *Arabidopsis thaliana*, to test current assumptions on functional redundancy and trait robustness. Our analyses of *BRZ/BEH* gene mutants and mutant combinations revealed that functional redundancy among gene family members does not contribute to trait robustness. Connectivity is another commonly cited determinant of robustness; however, we found no correlation between connectivity among gene family members or their connectivity with other transcription factors and effects on robustness. Instead, we found that only *BEH4*, the most ancient family member, modulated developmental robustness. We present evidence that regulatory cross-talk among gene family members is integrated by *BEH4* and promotes wild-type levels of developmental robustness. Further, the chaperone HSP90, a known determinant of developmental robustness, appears to act via BEH4 in maintaining robustness of embryonic stem length. In summary, we demonstrate that even among closely related transcription factors, trait robustness can arise through the activity of a single gene family member, challenging common assumptions about the molecular underpinnings of robustness.

## INTRODUCTION

Development relies on the coordinated action of low concentrations of regulatory factors diffusing within and between cells, which inevitably results in random developmental errors. Typically, organisms tolerate developmental errors, resulting in canalized, wild-type-like individuals (Waddington 1942; Masel and Siegal 2009; Lempe *et al.* 2012; Whitacre 2012; Félix and Barkoulas 2015). Robustness to developmental errors is an intrinsic property of all organisms and is genetically controlled (Hall *et al.* 2007; Ansel *et al.* 2008; Sangster *et al.* 2008a; Rinott *et al.* 2011; Jimenez-Gomez *et al.* 2011; Metzger *et al.* 2015; Ayroles *et al.* 2015). However, the molecular mechanisms that regulate developmental robustness are poorly understood, which is largely due to the technical obstacles of studying this phenomenon in complex, multicellular organisms.

Regulation of developmental robustness has been attributed to a handful of molecular mechanisms and features of gene regulatory networks (reviewed in (Masel and Siegal 2009; Lempe *et al.* 2012; Whitacre 2012; Félix and Barkoulas 2015; Lachowiec *et al.* 2015b)). In *Caenorhabditis elegans*, large-scale double mutant analysis identified several highly connected chromatin modifiers as positive regulators of developmental robustness (Lehner *et al.* 2006). In *Arabidopsis thaliana*, QTL mapping for regulators of developmental robustness found evidence that the pleiotropic genes *ERECTA* and *ELF3* regulate developmental robustness (Hall *et al.* 2007; Jimenez-Gomez *et al.* 2011); both genes are also highly connected in genetic networks. Nevertheless, these and other plant studies suggest that robustness modulators act in a trait-specific rather than global manner, presumably through epistasis with several specific partner genes.

The protein chaperone HSP90, known for its role in promoting genetic robustness (Rutherford and Lindquist 1998; Queitsch *et al.* 2002; Yeyati *et al.* 2007; Jarosz and Lindquist 2010; Rohner *et al.* 2013; Lachowiec *et al.* 2013, 2015a) also maintains developmental robustness in many organisms. For example, HSP90 perturbation across many isogenic plants results in vastly increased phenotypic diversity (Queitsch *et al.* 2002; Sangster *et al.* 2007, 2008a). Similarly, naturally low levels of HSP90 correlate with greater penetrance of mutations in isogenic worms (Burga *et al.* 2011; Casanueva *et al.* 2012). HSP90’s apparently global role in developmental robustness of plants and animals is consistent with the chaperone’s exceedingly high connectivity in genetic networks (*i.e.* epistasis with many different partner genes), particularly with many genes encoding kinases and transcription factors important for growth and development (Taipale *et al.* 2010; Lachowiec *et al.* 2015a).

Theoretical and empirical studies suggest that developmental robustness emerges from the circuitry of genetic networks. For example, highly connected nodes in genetic networks may be of particular importance in regulating robustness to noise due to their many interactions (Levy and Siegal 2008; Masel and Siegal 2009; Whitacre 2012). Another feature of genetic networks commonly associated with developmental robustness is functional redundancy among genes (Gutiérrez and Maere 2014). Functional redundancy will compensate for stochastic losses of function in specific gene family members or paralogs (DeLuna *et al.* 2008, 2010).

Gene duplication is one obvious source of network redundancy, and thereby developmental robustness. In *Arabidopsis thaliana*, one-third of genes belong to multi-member gene families (Swarbreck *et al.* 2008), which have arisen through three well-supported whole genome duplications (Simillion *et al.* 2002; Bowers *et al.* 2003), in addition to segmental and tandem duplication events (The Arabidopsis Initiative 2000). Duplication of transcription factor genes provides a plausible but potentially complex form of robustness regulation. Transcription factor family members recognize highly similar DNA motifs (Franco-Zorrilla *et al.* 2014) and often regulate one another (Phillips and Hoopes 2008), showing functional redundancy as well as feedback regulation (Wang *et al.* 2012; Sullivan *et al.* 2014; Lachowiec *et al.* 2015b). At the same time, transcription factors are particularly vulnerable nodes for developmental robustness due to their often low cellular concentrations and positions as both master regulators (Chan and Kyba 2013) and endpoints of signaling cascades (Li *et al.* 2014). It is unclear how these different features of transcription factors and their gene families converge to regulate developmental robustness.

The *<u>BE</u>SI/BZRI <u>H</u>OMOLOG*(*BEH)* transcription factors belong to a small gene family exclusive to plants. With only six members (Wang *et al.* 2002), this family is tractable for studying the role of redundancy, connectivity, and feedback on developmental robustness. The well-studied founding members of the *BEH* family, *<u>B</u>RII-<u>E</u>MS-<u>S</u>UPRESSOR<u>1</u>* (*BES1*) and *<u>B</u>RASSINA<u>Z</u>OLE-<u>R</u>ESISTANT<u>1</u> (BZR1*) result from the most recent whole genome duplication in the *A. thaliana* lineage and are highly similar in sequence (Blanc *et al.* 2003). They are thought to be the primary transcription factors in brassinosteroid signaling; studies of phenotypic effects are largely restricted to dominant mutants (Zhao *et al.* 2002; Yin *et al.* 2002; Wang *et al.* 2002). Brassinosteroid signaling regulates a large number of physiological processes in plants, ranging from seed maturation to senescence (Clouse 2002). Brassinosteroids are recognized by the membrane-associated receptor BRI1 that then represses the activity of the GSK3 kinase BIN2. In the absence of brassinosteroids, BIN2 phosphorylates and inhibits BES1 and BZR1 (Zhao *et al.* 2002). In this phosphorylated state, BES1 and BZR1 are prohibited from entering the nucleus (Gampala *et al.* 2007). In the presence of brassinosteroids, BES1 and BZR1 are dephosphorylated (Tang *et al.* 2011) and localize to the nucleus, where they activate and repress different sets of targets genes (Yin *et al.* 2005; He *et al.* 2005; Sun *et al.* 2010; Yu *et al.* 2011). BES1 and BZR1 are known to interact with several other proteins to regulate transcription. For example, BES1 dimerizes with BIM family proteins (Yin *et al.* 2005) to increase DNA binding affinity *in vitro*, interacts with its target gene MYBL2 (Ye *et al.* 2012), and works with ISW1 (Li *et al.* 2010a), ELF6, and REF6 (Yu *et al.* 2008) to alter chromatin accessibility. Some studies have revealed differences in BES1 and BZR1 protein interactions. For example, BES1, but not BZR1, interacts with the known robustness regulator HSP90 (Shigeta *et al.* 2013; Lachowiec *et al.* 2013).

In contrast, the other family members *BEH1-4* are little studied, largely due to the lack of well-characterized loss-of function or dominant mutants. As BES1 and BZR1, BEH1, BEH2, BEH3, and BEH4 are thought to act as transcription factors (Wang *et al.* 2002; He *et al.* 2005). Moreover, BEH1, BEH2, BEH3, and BEH4 are phosphorylated in a manner similar to BES1 and BZR1 (Yin *et al.* 2005), and yeast two-hybrid analyses show that BEH2, in addition to BES1 and BZR1, interacts with a GSK3 kinase (Rozhon *et al.* 2010). In sum, previous studies support that BEH1, BEH2, BEH3, and BEH4 act redundantly with the well-studied transcription factors BES1 and BZR1 (Krizek 2009; Ye *et al.* 2012).

Here, we systematically examined the entire *BEH* family for effects on developmental robustness through the lenses of redundancy, connectivity, and feedback. Contrary to commonly held assumptions about the importance of redundancy and connectivity in robustness, we observe that robustness in hypocotyl growth arises largely due to the function of a single gene, *BEH4*, which appears to maintain proper cross-talk among *BEH* family members. Further, we trace HSP90’s role in maintaining robustness of hypocotyl length to the function of *BEH4*, thereby elucidating how this well-known regulator of global developmental robustness specifically affects this trait.

## METHODS

### Plant materials and growth conditions

*bes1-2* (Lachowiec *et al.* 2013), *bzr1-2* (GABI_857E04), *beh3-1* (SALK_017577), and *beh4-1* (SAIL_750_F08) are in the Col-0 background. *beh1-1* (SAIL_40_D04) and *beh2-1* (SAIL_76_B06) are in the Col-3 background. Using qPCR (see below), we confirmed that none of the mutants produced full-length transcripts; most produced no transcript at all.

For hypocotyl length assays, seeds were sterilized for 10 minutes in 70% ethanol, 0.01% Triton X-100, followed by 5 minutes of 95% ethanol. After sterilization, seeds were suspended in 0.1% agarose and spotted on plates containing 0.5x Murashige Minimal Organics Medium and 0.8% bactoagar. Seeds on plates were then stratified in the dark at 4°C for 3 days and then transferred to an incubator cycling between 22° for 16 hours and 20° for 8 hours to imitate long days. Plate position was changed every 24 h to minimize position effect for light grown seedlings. Racks of plates containing dark-grown seedlings were wrapped in foil. For HSP90-inhibitor assays, 1μM geldanamycin (Sigma) was suspended in the medium. Equivalent amounts of the solvent DMSO were used for control treatment.

### Phenotyping

For estimates of hypocotyl CV, three replicates of n > 50 were measured. Assays of mean hypocotyl length were completed in triplicate with n > 15. Photos were taken of each plate, and individual hypocotyls were manually measured using NIH ImageJ1.46r.

### qPCR

Three biological replicates of sixty pooled 5-day dark grown seedlings were harvested. Tissue was frozen in liquid nitrogen and ground by hand with a pestle. RNA was extracted using the SV Total RNA Isolation kit (Promega). To remove contaminating DNA, a second DNase treatment was completed according to the Turbo DNase protocol (Ambion). Poly-A tail cDNA was produced using LightCycler kit with oligo-dT primers (Life Technologies). qPCR primers are listed in Table S3. In both the *bzr1-2* and *beh2-1* mutants these qPCR primers amplified products. The absence of the full-length transcripts was confirmed using primers that target the full-length transcript.

## RESULTS

### *BEH* family members share function in regulating hypocotyl elongation in the dark

To dissect the individual functions of different members of the *BEH* family, equivalent mutants, ideally recessive, complete loss-of-function (*lof*) mutants, are required for genetic analysis. For studies of *BES1* and *BZR*, researchers have largely relied on the dominant mutants *bes1-D* and *bzr1-1D*, which introduce the same nucleotide change in their respective PEST domains (Yin *et al.* 2002; Wang *et al.* 2002). This mutation appears to stabilize PEST interaction with a de-phosphatase PP2A (Tang *et al.* 2011), thereby creating dominant mutants that are constitutively active. Not all members of the *BEH* family are predicted to contain homologous PEST domains (Rogers *et al.* 1986) (Figure S1), so comparable dominant mutants cannot be created. To generate comparable *lof* mutants, we acquired T-DNA insertion mutants for each gene family member (*bes1-2, bzr1-2, beh1-1, beh2-1, beh3-1*, and *beh4-1*) (Lamesch *et al.* 2012) (Figure S1). Based on expression analysis, we are confident that we have generated complete *lof* mutants for each family member, thereby enabling unbiased phenotype comparisons.

The phenotypes of *bes1-D* and *bzr1-1D* included hyper-elongation of hypocotyls when grown in the dark (Yin *et al.* 2002; Wang *et al.* 2002), suggesting that BEH1, BEH2, BEH3, and BEH4 may function in promoting hypocotyl growth. Indeed, the *bes1-2, bzr1-2, beh3-1*, and *beh4-1* recessive *lof* mutants produced significantly shorter hypocotyls than wild-type in the dark (Figure 1a, p < 0.0001, linear mixed effects model, n = 70), demonstrating that these four family members, but not BEH1 and BEH2, are positive regulators of dark growth. Our results are consistent with previous findings in which RNAi targeting *BES1* reduces hypocotyl length (Yin *et al.* 2005; Wang *et al.* 2013), and the *bes1-1* T-DNA insertion mutant exhibits reduced hypocotyl length (He *et al.* 2005). Curiously, the recessive *lof* mutants of the founding and best-studied members of the *BEH* family, *BES1* and *BZR1*, were not the most affected in dark growth; the *lof* mutants of the ancestral members *BEH3* and *BEH4* showed larger effects on dark growth, with *beh4-1* exhibiting the strongest defect (Figure 1a), though the effect size was still small. The small but significant effects in these four mutants suggest that these gene family members share function but are not fully redundant in regulation hypocotyl growth in the dark.

**Figure 1.**
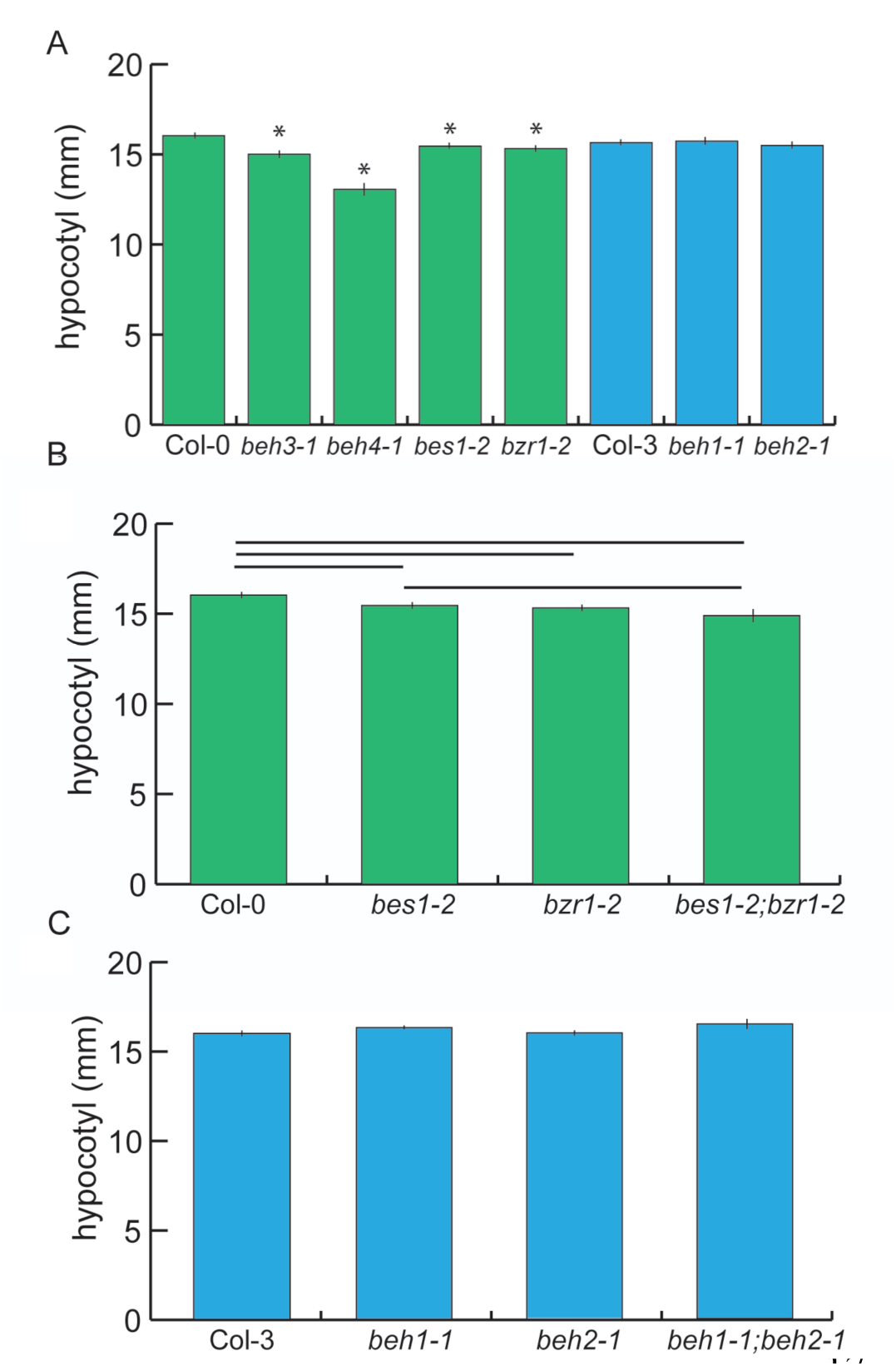
The *BEH* family encodes genes with non-redundant effects on hypocotyl length. **A)** Seedlings were grown for seven days in the dark, and hypocotyls were measured. *beh3-1, beh4-1, bes1-2, bzr1-2* hypocotyls were significantly shorter than those of wild-type (* p < 0.0001, linear mixed effects model with genotype as a fixed effect and replicate as a random effect). **B)** The phenotype of the *bes1-2;bzr1-2* double mutant suggests that *BZR1* is epistatic to *BES1* because there was no significant difference in hypocotyl length between *bes1-2* and *bes1-2;bzr1-2.* Significant differences (p < 0.05) are displayed by the horizontal bars as determined by linear mixed effect modeling. **C)** No significant differences in hypocotyl length were observed for *beh1-1* and *beh2-1* single mutants, or for the double mutant *beh1-1;beh2-1*. For **A-C)** one representative replicate experiment with standard error of the mean for n > 20 is shown.

There was no significant difference in dark growth between *bes1-2* and *bzr1*-*2* mutant seedlings, suggesting that *BES1* and *BZR1* contribute to dark growth to the same degree (Figure 1a). This finding is consistent with the similar phenotypes of the dominant *bes1-D* and *bzr1-1D* mutants (Lachowiec *et al.* 2013); it is also consistent with the high sequence identity between *BES1* and *BZR1* (Wang *et al.* 2002), and their overlapping patterns of expression (Yin *et al.* 2002; Wang *et al.* 2002). To determine whether BES1 and BZR1 independently (*i.e.* additively) regulate dark growth, we examined the *bes1-2;bzr1-2* double mutant. The double mutant tended to be shorter than either single mutant, but was only significantly shorter than *bes1-2* (Figure 1b), suggesting that *BZR1* is epistatic to *BES1* in promoting hypocotyl growth in the dark. Thus, although *BZR1* and *BES1* do not act fully redundantly in hypocotyl elongation, they appear to have overlapping rather than independent functions in its regulation. We speculate that these degenerate functions of *BES1* and *BZR1* may arise from different interacting protein partners.

Among the six family members, *BEH1* and *BEH2* did not affect dark growth (Figure 1a). Both genes are highly similar in sequence. To explore potential functional redundancy between *BEH1* and *BEH2*, we created the respective double mutant and assessed hypocotyl growth. The double mutant *beh1-1;beh2-1* exhibited no significant growth defect compared to wild-type, or the single mutants *beh1-1*, or *beh2-1* (Figure 1c). This result indicates either that *BEH1* and *BEH2* do not regulate hypocotyl elongation or that they act redundantly with other family members or other, unrelated genes in regulating dark growth. Taken together, BES1, BZR1, BEH3, and BEH4 share the function of regulating hypocotyl growth during growth in the dark. Notably, the *beh4-1* single mutant was significantly shorter than the *bes1-2;bzr1-2* double mutant (p < 0.0001, linear mixed effects model, n = 70), demonstrating *BEH4’s* dominant role in controlling dark growth.

In addition to skotomorphogenesis, *BES1* and *BZR1* are also important for photomorphogenesis and flowering (Li *et al.* 2010b) based on *bes1-D* and *bzr1-1D* phenotypes. BES1 is known to interact with the flowering time regulating proteins, ELF6 and REF6 (Yu *et al.* 2008). We detected no significant defects for the *BEH* family recessive *lof* mutants for flowering time (Figure S2), which agrees with earlier findings for a *BES1* T-DNA insertion line, *bes1-1* (He *et al.* 2005).

When grown in the light, *bes1-D* and *bzr1-1D* exhibit opposing effects on hypocotyl growth, with *bzr1-1D* showing shortened hypocotyls (He *et al.* 2005; Gampala *et al.* 2007). In previous work, the recessive line *bes1-1* showed reduced growth in the light (He *et al.* 2005). Therefore, we examined all of our recessive mutants for light growth. As light-grown seedlings have very short hypocotyls, at least 70 seedlings per genotype were required to detect significant differences for an effect size of 0.5mm (power analysis, power = 0.8). We hypothesized that the *bzr1-2* would show longer hypocotyls than wild-type in the light, based on the shortened *bzr1-1D* phenotype. Indeed, *bzr1-2* showed significantly longer hypocotyls than wild-type (p = 0.0087, linear mixed effects model, n = 70, Figure S3). In summary, our results reveal extensive, yet not complete, functional redundancy among these closely related transcription factors and emphasize the importance of using recessive, *lof* mutants for genetic analysis to elucidate function and primacy of individual genes in gene families

### *BEH4* is a determinant of robustness

We hypothesized that the observed extensive functional redundancy among BES1, BZR1, BEH3, and BEH4 may contribute to developmental robustness of dark grown hypocotyls (Wagner 2000; Gu *et al.* 2003; Lachowiec *et al.* 2015b). Measuring developmental robustness is straightforward in isogenic lines. By growing isogenic lines randomized in the same controlled environment, any variation in phenotype is attributed to errors in development and used as a measure of developmental robustness (Waddington 1942; Queitsch *et al.* 2002). Developmental robustness is often expressed as the coefficient of variation or CV (s^2^/u) (Lempe *et al.* 2012; Geiler-Samerotte *et al.* 2013; Gutiérrez and Maere 2014). We measured hypocotyl length with high replication in the *BEH* family single mutants using a randomized design to control for micro-environmental differences. Mutants in the founding members of the *BEH* family, *bes1-2* and *bzr1-1* did not significantly affect developmental robustness in hypocotyl length. Instead, *beh4-1* showed a highly replicable and significant decrease in developmental robustness (Figure 2a, p = 3.145* 10^−7^, Levene’s test, n = 210). No other single mutant significantly affected robustness. We conclude that robustness in dark grown hypocotyls was most affected by *BEH4* activity, which also affected trait mean the most (Figure 1a). This result recalls the results of a prior study, in which we found that HSP90-dependent loci for developmental robustness of dark grown hypocotyls often coincide with those for trait means upon HSP90 perturbation (Sangster *et al.* 2008a).

**Figure 2.**
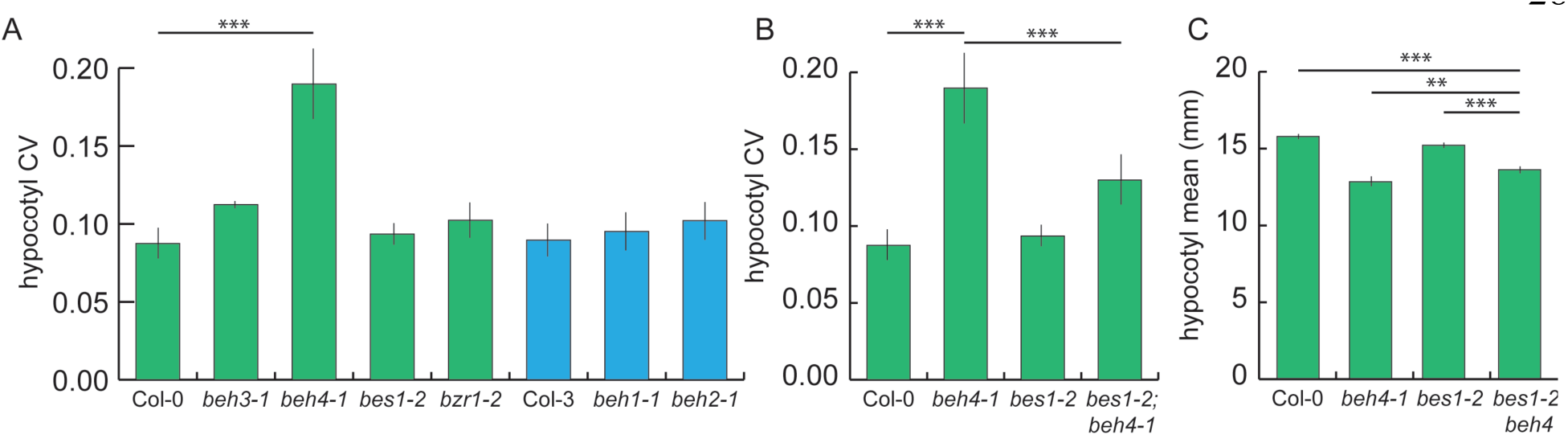
*BEH4* contributes the most to robustness of dark grown hypocotyls. **A)** The *beh4-1* mutant exhibits significantly greater variation in hypocotyl length compared to wild-type (*** p < 0.0001, Levene’s test, n = 210). None of the other single mutants increase hypocotyl length variance significantly. **B)** The double mutant *bes1-2;beh4-1* showed an intermediate effect on hypocotyl length robustness compared to either single mutant (*** p < 0.0001, Levene’s test, n = 210). CV was estimated in three biological replicates. Standard error of the mean for n = 3 is shown for both **A)** and **B)**. **C)** The double mutant *bes1-2;beh4-1* also showed an intermediate effect on hypocotyl mean values compared to either single mutant (*** p < 0.0001, ** p < 0.001, linear mixed effects model with genotype as a fixed effect and replicate as a random effect).

We hypothesized that we may observe a further loss of robustness by removing additional functional *BEH* family members. We examined the *bes1-2;beh4-1* double mutant because both single mutants affected mean hypocotyl length in the dark. Surprisingly, we found that loss of BES1 activity partially rescued developmental robustness in the double mutant (Figure 2b). Similarly, loss of BES1 activity partially rescues mean trait value in the *bes1-2;beh4-1* compared to the single *beh4-1* mutant (Figure 2c). We conclude that BEH4 and BES1 do not act redundantly in generating hypocotyl developmental robustness and trait means. Rather, we suggest that developmental robustness arises through the integrated activity of various family members. Note that *bes1-2* alone did not affect developmental robustness. It is only through its interaction with *BEH4* that we observed its apparently stabilizing effect. Indeed, others have argued that BES1 directly or indirectly regulates *BEH4* as shown by ChIP-analysis, at least in aerial tissues (Yu *et al.* 2011).

### Expression feedback among members of the *BEH* family in the light and dark

We wanted to further explore the putative functional integration among *BEH* family members that may underlie *BEH4*-dependent developmental robustness. Specifically, we hypothesized that *BEH4* acts as hub gene among *BEH* family members. Highly connected hub genes such as the well-characterized HSP90 are thought to affect robustness through their interaction with many other loci; hub perturbation results in large-scale phenotypic effects and loss of robustness (Levy and Siegal 2008; Fu *et al.* 2009; Lempe *et al.* 2012; Lachowiec *et al.* 2015b). BES1 and BZR1 ChIP results (Sun *et al.* 2010; Yu *et al.* 2011) suggest that all other *BEH* family members are potential transcriptional targets of BES1 and BZR1 (Table S1), consistent with direct or indirect regulation among family members. Further, expression of *BEH2* is up-regulated in RNAi lines in which *BES1* is targeted (Wang *et al.* 2013), and *BZR1* expression is reduced in *bes1-1* mutants (Jeong *et al.* 2015). To test our hypothesis that BEH4 is the most highly connected genes in this gene family, we determined the relative expression of each *BEH* family member in each single *lof* mutant background. If mean gene expression was altered more than 2-fold in a given mutant background, we assumed a direct or indirect genetic interaction between the assayed and the mutated gene. Disproving our hypothesis, we found that *BEH3* was the most highly connected gene among the *BEH* family, not *BEH4* (Figure 3). Seven connections among *BEH3* and other family members were counted, with *BEH3* directly or indirectly regulating three family members and *BEH3* expression affected in four mutants. Two of these interactions were reciprocal, in which *BEH3* and *BEH4* regulate each other, as well as *BEH3* and *BES1*. Similar to *BEH3, BEH4* directly or indirectly affected gene expression of three family members, but only two mutants influenced *BEH4* expression. Notably, the *lof beh3-1* mutant showed no decrease in developmental robustness; hence, connectivity, another frequently cited cause of developmental robustness (Levy and Siegal 2008; Lachowiec *et al.* 2015b), is apparently not majorly involved in robustness of hypocotyl growth. This interpretation does not consider possible interactions at the protein level through heterodimers among family members or connections of *BEH4* with genes outside its gene family.

**Figure 3.**
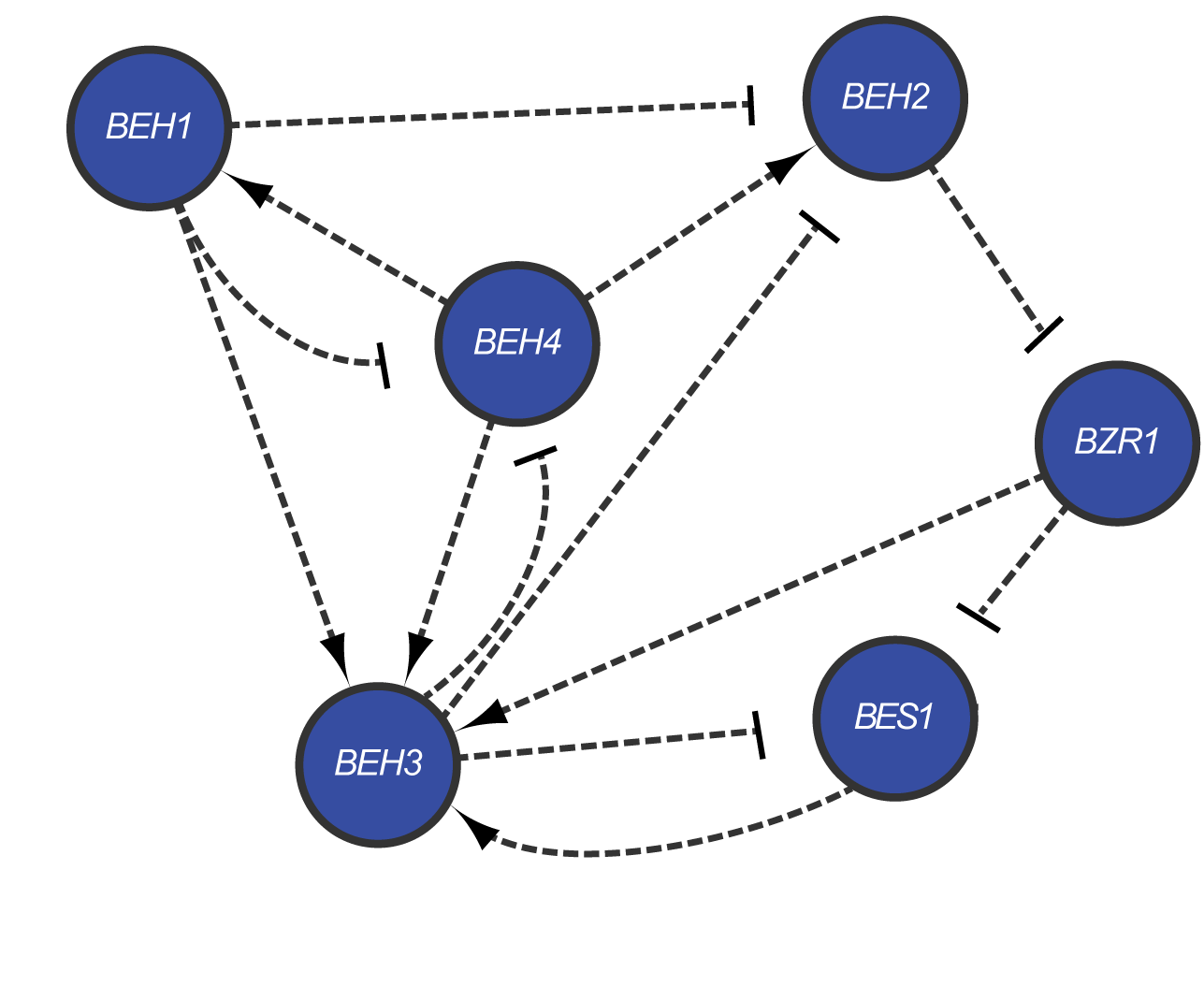
*BEH* family members engage in extensive regulatory cross-talk. Direct or indirect regulatory relationships among *BEH* family members were determined using qPCR. A regulatory relationship was called for a gene if a greater than a 2-fold expression difference between wild-type and mutant backgrounds was measured. Both positive and negative regulatory relationships are indicated.

Although connectivity was not associated with phenotypic effects, gene duplicate age appeared to be associated with the number of connections among family members. *BES1* and *BZR1* are the most recently duplicated members of the family, followed by *BEH1* and *BEH2*, with *BEH3* and *BEH4* being the most ancestral (Blanc *et al.* 2003). With three connections, BZR1 and *BES1* were the least connected genes; *BEH1* and *BEH2* each showed four direct or indirect regulatory connections. These results are consistent with closely related transcription factors gaining regulatory complexity over time as paralogs are added.

To further explore the regulatory network underlying hypocotyl elongation in the dark, we analyzed recent DNaseI-seq data of dark grown seedlings (Sullivan *et al.* 2014). We and others have suggested that robustness regulators may be characterized by numerous regulatory inputs and few outputs, an architecture well suited to buffer noise (Sangster *et al.* 2004; Lehner *et al.* 2006; Levy and Siegal 2008; Rinott *et al.* 2011). Therefore, we identified and counted transcription factor (TF) binding motifs in the accessible chromatin marking the putative promoters of all *BEH* gene family members (Table S2). The promoter-proximal accessible chromatin of *BEH4* and *BEH3* each contained 25 TF binding motifs; 26 TF motifs were found for *BEH2* and 35 for *BZR1*. In contrast, no TF motifs were detected for *BEH1*, and only six TF motifs were found for *BES1*. We conclude that at least for the BEH gene family the number of regulatory inputs (measured as number of promoter TF binding sites) is not associated with the severity of phenotypic effects on developmental robustness or trait mean. We were unable to assess regulatory outputs because the binding motifs of individual *BEH* family members are unknown. *BES1* and *BZR1* both recognize the BRZ motif, which resided in accessible, promoter-proximal chromatin of 230 genes. Although *BEH4* most strongly affects phenotype among the BEH family members, neither connectivity nor regulatory architecture is consistent with the hypothesized role of BEH4 as a hub gene.

### HSP90 likely maintains developmental robustness of dark-grown hypocotyls via BEH4

HSP90 function is crucial for developmental robustness of dark-grown hypocotyls and other traits (Queitsch *et al.* 2002; Sangster *et al.* 2007, 2008a; b). As HSP90 chaperones the *BEH* family member BES1 (Shigeta *et al.* 2013; Lachowiec *et al.* 2013), we hypothesized that the dominant role of *BEH4* in developmental robustness may involve HSP90. To test this hypothesis, we assessed the genetic interaction of HSP90 and *BEH4*, using the potent and highly specific inhibitor geldanamycin (GdA) to reduce HSP90 function. As previously observed, HSP90 inhibition in wild-type seedlings decreased robustness (Figure 4a). HSP90 inhibition in *bes1-2* mutant seedlings also decreased robustness, closely resembling the phenotypic effect observed in wild-type (Figure 4a). In stark contrast, *beh4-1* exhibited no change in developmental robustness upon HSP90 inhibition (p=0.296, Levene’s test, n=210). In fact, *BEH4* appeared to be epistatic to HSP90 in mediating developmental robustness of dark-grown hypocotyls, suggesting that HSP90 acts via BEH4 in this pathway.

**Figure 4.**
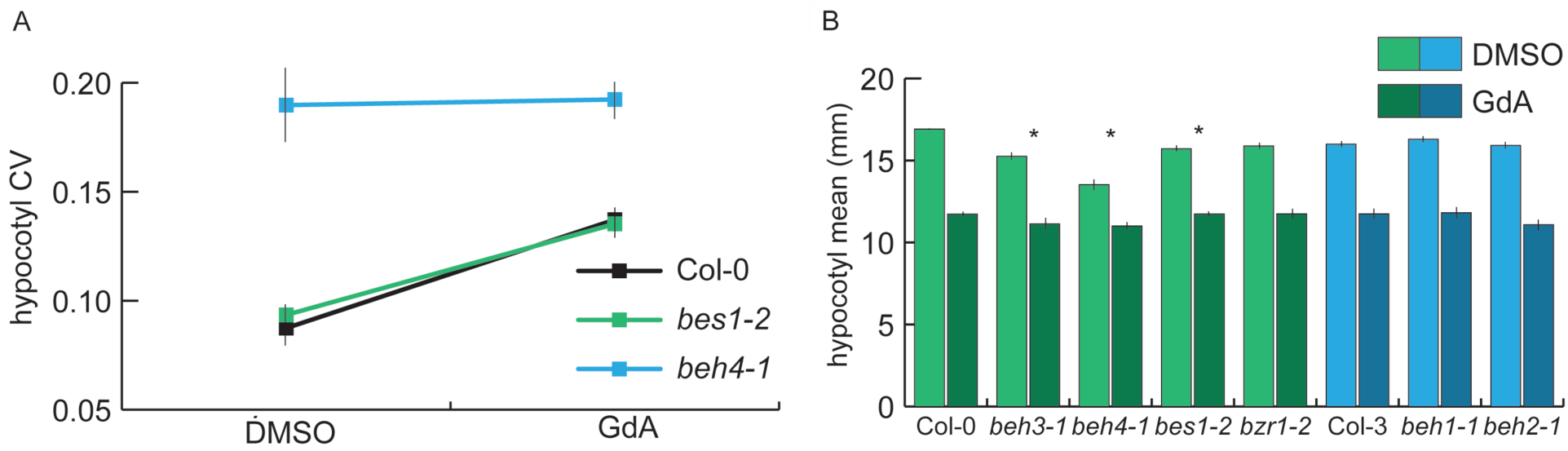
Robustness provided by HSP90 likely arises from the chaperone’s interaction with BEH4. **A**) Seedlings were grown with or without HSP90, and hypocotyl length was measured in three replicate experiments. CV was calculated for each replicate and the standard errors of the mean for n = 3 are shown. BES1 is a known HSP90 client in this gene family. **B)** Hypocotyl length mean data for the same conditions are shown. One representative replicate experiment with standard error of the mean for n > 20 is shown. * Significant differences in mean trait response to HSP90 inhibition are shown (p < 0.03, linear mixed model with genotype, treatment, and interaction effects as fixed effects and replicate as a random effect).

The most obvious mechanism by which HSP90 would act via BEH4 to mediate developmental robustness is by chaperoning BEH4. The BEH family member BES1, but not BZR1, is an HSP90 client (Shigeta *et al.* 2013; Lachowiec *et al.* 2013). Due to the high similarity among BEH family members, it is certainly likely that others are also HSP90 substrates, as client status is often shared among family members (Taipale *et al.* 2012; Lachowiec *et al.* 2015a). HSP90 inhibition typically compromises the function of its clients due to mis-folding and degradation (Taipale *et al.* 2010). The observed epistasis of BEH4 with HSP90 in developmental robustness (lack of response in *beh4-1* upon HSP90 inhibition) is consistent with the hypothesis that BEH4 is an HSP90 client.

To further test this hypothesis, we analyzed trait means of all single mutants of the *BEH* family members with and without HSP90 inhibition. As expected from our previous studies (Lachowiec *et al.* 2013), the *lof* mutant of the HSP90 client BES1, *bes1-2*, was significantly less sensitive than wild-type to HSP90 inhibition (p = 0.03, linear mixed effects model, n = 20, Figure 4b). Moreover, both *beh3-1*, and *beh4-1* were significantly less affected than wild type (p = 0.01, p < 0.0001, respectively, linear mixed effects model, n = 20, Figure 4b). In contrast, BRZ1, which is not chaperoned by HSP90 (Shigeta *et al.* 2013; Lachowiec *et al.* 2013), BEH1 and BEH2 behave like wild type. These results are consistent with our hypothesis that BEH4 and possibly BEH3 are HSP90 clients.

## DISCUSSION

Developmental robustness is thought to emerge from the topology of gene networks, including the activity of redundant genes, gene connectivity, and regulatory architecture (Lachowiec *et al.* 2015b). Here, we trace robustness of the model trait hypocotyl length to a specific member of the *BEH* gene family, *BEH4*. Contrary to our expectations, *BEH4*’s role in developmental robustness of dark-grown hypocotyls does not appear to arise through functional redundancy with closely related family members. Loss of another family member did not further decrease developmental robustness; rather, we observed partial rescue. *BEH4*, the ancestral member of the *BEH* family, also showed the largest effect on the trait mean phenotype. Our observations challenge a prior theory that additional connections (here paralogs), added later, may stabilize traits (Wagner 1996). Instead, at least for this particular trait and gene family, the ancestral gene remained the largest player for both trait mean and variance (developmental robustness). Previous studies frequently found that loci that affect trait robustness also affect trait mean (Hall *et al.* 2007; Sangster *et al.* 2008a; Ordas *et al.* 2008; Jimenez-Gomez *et al.* 2011). This frequently observed overlap makes intuitive sense: a gene that significantly affects trait mean when disrupted will perturb the underlying stabilizing genetic network and may so decrease trait robustness (Félix and Barkoulas 2015). As stabilizing selection on genetic variants that affect both mean and variance will be far stronger than selection on variants that affect only trait variance, genes such as *BEH4* will play critical roles in maintaining phenotypic robustness.

Gene network hubs are thought to be crucial for developmental robustness, presumably due to their high number of connections with other loci. This assumption is certainly supported by several prior studies in plants, yeast and worms (Queitsch *et al.* 2002; Lehner *et al.* 2006; Sangster *et al.* 2007; Levy and Siegal 2008; Rinott *et al.* 2011). At the small scale of the *BEH* gene family this assumption did not hold true. We did, however, observe that the older gene duplicates, *BEH3* and *BEH4*, tended to engage in more regulatory connections than other family members, consistent with previous studies finding that number of protein interactions correlates with gene age (Eisenberg and Levanon 2003; Kunin *et al.* 2004; Saeed and Deane 2006). However, *beh3-1* did not exhibit altered developmental robustness, indicating that connectivity alone does not suffice to explain effects on developmental robustness.

One may argue that our experiments did not thoroughly test *BEH4* as a hub, as we primarily restricted our analysis to the *BEH* family. The known genetic network underlying hypocotyl dark growth is certainly complex (Oh *et al.* 2014), and thus far *BEH4*’s role within this network has been unknown. Our analysis of DNAseI-seq data for dark-grown seedlings revealed the putative number of TFs regulating different *BEH* family members (Table S1). The number of potential regulatory inputs for individual family members did not correlate with the severity of the phenotypic effects in their mutants; several family members showed equal or more inputs that *BEH4*.

Our data best support the alternative hypothesis that *BEH4*’s role in developmental robustness arises through the topology of its connections with other family members. For example, feedback loops are known to promote robustness (Hornstein and Shomron 2006; Ebert and Sharp 2012; Cassidy *et al.* 2013; Lachowiec *et al.* 2015b). We found that *BEH4* positively regulates *BEH3* and *BEH1*, which in turn, both negatively regulate *BEH4*. Hence, loss of robustness in *beh4* mutants likely arises through the loss of finely tuned regulation among family members. This hypothesis is supported by our observation that in the *bes1-2*;*beh4-1* double mutant developmental robustness is partially rescued, possibly because the fine-tuned balance among family members is partially restored in the double mutant.

The BEH family member BES1 is known to be a client of the developmental robustness regulator HSP90 (Shigeta *et al.* 2013; Lachowiec *et al.* 2013). HSP90 presumably governs developmental robustness by chaperoning its client proteins, which function in diverse developmental pathways (Taipale *et al.* 2010). HSP90 inhibition leads to destabilization and loss of function for its many clients (Xu 1993; Taipale *et al.* 2012). Notably, loss of BES1 function did not affect robustness, indicating that HSP90 does not regulate robustness through its client BES1. Instead, we observed that HSP90-dependent robustness of hypocotyl growth is likely due to *BEH4* function—unlike wild type, the *beh4-1* mutant showed no response to HSP90 inhibition with regard to developmental robustness. Together, this result and the significantly diminished mean response of *beh4-1* mutant to HSP90 suggest that BEH4 is also an HSP90 client. In sum, we propose that HSP90 regulates developmental robustness of dark-grown hypocotyls through the activity of BEH4, which is central for fine-tuned cross-regulation among all *BEH* family members.

## ACKNOWLEDGMENTS

We thank Alessandra Sullivan for sharing *BEH* family DNaseI-seq results. This work was supported by grants from the National Human Genome Research Institute Interdisciplinary Training in Genomic Sciences (T32 HG00035 to J.L. and G.A.M.), the National Science Foundation (DGE-0718124 to J.L. and DGE–1256082 to G.A.M.) and National Institutes of Health (new innovator award no. DP2OD008371 to C.Q.).

